# Artificial Intelligence-Driven Nanosensing to Identify Circulating Bacterial DNA: A Novel Approach to Breast Cancer Risk Profiling

**DOI:** 10.64898/2026.09.28.755235

**Authors:** Nazim Nazeer, Pooja Ratre, Vikas Gurjar, Olga Goryacheva, Santram Lodhi, Arpit Bhargava, Irina Yu Goryacheva, Pradyumna Kumar Mishra

## Abstract

Perovskite quantum dots (PQDs) are exceptionally promising next-generation optical materials for bioimaging applications. This paper presents an optical PQDs nanocomposite method to identify bacterial species associated with breast cancer, employing a LSTM deep learning model. The results indicate that the developed PQD-based nano-sensor offers impressive sensitivity, selectivity, and practical application. The four bacterial species identified in the study, *Pseudomonas aeruginosa, Vibrio parvula, Acinetobacter baumannii*, and *Streptococcus vestibularis*, were considered the most relevant indicators of breast cancer. With rapid detection, an intuitive design, and improved reliability, the PQD sensor could be an excellent breast cancer diagnostic tool for a point-of-care application. Furthermore, it has the potential to be integrated into healthcare systems, specifically in resource-limited settings, thereby improving the accessibility and efficiency of early breast cancer detection.

## 1. Introduction

The human microbiome is essential to various human physiology including digestion, immune system modulation, protection against pathogenic organisms, and synthesis of certain vitamins, with most microorganisms believed to be nontoxic or useful. Some microbes have been identified as harmful to human health, particularly those associated with malignancies and other conditions marked by abnormal inflammation [1]. Dysbiosis, characterized by a microbial imbalance where pathogenic bacteria surpass beneficial strains, may result in several types of disease including breast cancer (BC) [2]. The inhabitants of the gastrointestinal system are proving to be significant regulators of breast carcinogenesis. Breast tissue hosts its own microbiome, which may influence cancer risk and progression [3-4]. The microbiome influences BC by affecting immune responses, inflammation, and estrogen metabolism. The gut microbiota, through the estrobolome, reactivates estrogens, raising their levels and promoting estrogen receptor-positive (ER+) BC growth. Dysbiosis disrupts this balance, prolonging estrogen exposure, a risk factor for BC. Moreover, CYP19A1 gene is also involved in the estrogen synthesise, that also supports the ER+ BC Development. Similarly, estrogen is also modulated by Gut microbiota mainly in women with postmenopausal [5, 2]. Inflammatory cytokines like interleukin-6 (IL-6) and tumor necrosis factor-alpha (TNF-α), driven by microbial changes, can activate the STAT3 pathway, contributing to BC proliferation and metastasis. The gut microbiome also plays a role in regulating genes associated with DNA repair mechanisms and cellular homeostasis, including *BRCA1* and *BRCA2*, which are crucial for maintaining genomic integrity [6-7]. However, recent studies suggested that microbial dysbiosis could also contribute to sporadic BC cases by affecting DNA repair processes. Additionally, the microbiome can influence epigenetic modifications, including DNA methylation and histone acetylation, which are critical regulators of gene expression in cancer-related pathways. For instance, abnormal methylation of tumor suppressor genes like *TP53* can occur due to chronic inflammation induced by microbial imbalance, contributing to uncontrolled cell growth and BC progression [8]. Recent advancements in 16S rRNA gene sequencing have enabled researchers to detect changes in the gut and breast microbiota. This results in the identification of bacterial species associated with immune modulation and provides insights into microbial imbalances influencing BC development [9-11].

Perovskite quantum dots (PQDs), a class of quantum dots (QDs) based on perovskite materials, have garnered significant attention for their potential in various electronic and optoelectronic applications due to their impressive properties. These nanocrystals, typically ranging from 2-10 nanometers in diameter, exhibit unique quantum-size effects, offering tunable and efficient photoluminescence (PL) and narrow emission spectra [12]. Perovskites, with their versatile ABX3 crystal structure, allow for the incorporation of various cations, enabling the development of diverse materials with interesting properties such as superconductivity and ferroelectricity. When perovskite materials are used to create PQDs, they can surpass traditional metal chalcogenide QDs, particularly in their tolerance to defects, high photoluminescence quantum yields, and superior colour purity [13].

Previously, an advanced sensing technique for the detection of 16s rRNA of cardiovascular diseases related bacterial species has been published [14]. Building on this foundation, this study exploits the unique properties of PQDs, including their photoluminescence and high stability, to enhance the sensitivity and specificity of 16S rRNA detection in BC, providing a more robust platform for early diagnostics. In the current work we have identified different bacterial species in the circulating microbiome that are associated with BC, with some species playing a more significant role in disease progression than others. To systematically analyse these findings, we developed an artificial intelligence (AI) model that examine large microbiome datasets. This model enabled us to identify the most abundant and most strongly BC-associated bacterial species. We found *A. baumanii S. vestibulacis, P. aeruginosa and V. parvula* are involved in the mechanism of the BC development [15]. Detecting 16S rRNA from these species could enhance early detection. In our current work, we developed PQD-based nanobiosensors for sensitive 16S rRNA detection. Our detection strategy utilizes streptavidin-conjugated PQDs that bind to biotinylated probes, allowing for specific and integrated detection of 16S rRNA from these target bacteria. This combined approach not only enhances early diagnostic potential but also provides deeper insight into the complex microbial interactions influencing breast cancer development [16-17]

## 2. Material and Methods

### 2.1. Reagents

PQDs were used from Merck Sigma-Aldrich, Inc. in St. Louis, MO, USA. NHS (N-hydroxysuccinimide) and EDC (1-ethyl-3-[3-dimethylaminopropyl] carbodiimide hydrochloride) were from the Thermo Fisher Scientific in Waltham, Massachusetts, USA. Pierce™ RNA 3’ End Biotinylation Kit was bought from the USA based company, Thermo Fisher Scientific in Waltham, MA. PCR (polymerase chain reaction) primers and the biotinylation probe was bought from the Eurofins Genomics in Anzinger Strasse, Germany and Integrated DNA Technologies in Coralville, Iowa, USA. The PBS solution (10X) was procured from the Himedia Laboratory in Mumbai, MH, India. The circulating nucleic acid isolation kit (QIAMP) was bought from the QIAGEN in Hilden, Germany. SYBR Safe and Master Mix were obtained from Thermo Fisher Scientific Waltham, MA, USA. The gel loading dye (6X) was purchased from the Himedia Laboratory (Mumbai, MH, India). Lastly, PrimeScript 1st strand cDNA synthesis kit was got from the Takara Bio Inc. in Shiga, Japan.

### 2.2. Artificial intelligence model for the prediction of the most abundant bacterial species

#### Blast dataset

The dataset used in this work, which focused on the metagenomic analysis of BC tissue, was obtained from the NCBI Sequence Read Archive (SRA) under project accession PRJNA926328. SRR23219779.fasta (Run accession: SRX19167651), SRR23219776.fasta (Run accession: SRX19167654), and SRR23219777.fasta (Run accession: SRX19167653) are the particular datasets that were utilised. To create a complete dataset for BLAST analysis, FASTA files were combined. To find and describe bacterial DNA fragments, BLAST was applied to the combined dataset. Using the makeblastdb program, a reference sequence database was first created, making sure it was structured correctly for nucleotide sequence comparison. In order to maximise computing efficiency during BLAST processing, the combined dataset was then divided into smaller halves, each of which had around 1,000 sequences. BLASTn was used to process each chunk, matching the query sequences to the reference database. Critical alignment data, including query and subject sequence IDs, percent identity, alignment length, mismatches, gaps, e-values, and bit scores, were provided in the tabular results (outfmt 6). For further investigation, the separate BLAST results from every piece were combined into a single file. A thorough investigation of the bacterial sequences found in breast cancer tissue samples was made possible by the effective processing and analysis of large-scale metagenomic data allowed feasible by this structured technique.

#### LSTM model development

In the next stage, we created an LSTM model to forecast which bacterial species were most common in the metagenomic data. Important numerical features like length (alignment length), pident (percentage identity), mismatch (number of mismatches), qstart (query start position), sstart (subject start position), gapopen (number of gap openings), qend (query end position), send (subject end position), evalue (expectation value), and bitscore were converted to numeric values as part of the preprocessing of the dataset, which was the first step in the methodology. To ensure the consistency and predictability of the data, non-numeric entries were forced to NaN and then eliminated. Label encoding was used to convert categorical data into a format that was appropriate for model training by encoding the sseqid column, which represents bacterial species identifiers, into numerical categories [18]. Then, in order provide a consistent contribution to the model training process and to minimise any potential biases driven on by different scales, features were normalised using the MinMaxScaler, scaling the values to range between 0 and 1. The information was transformed into the [samples, time steps, features] format needed for LSTM models. 80% of the dataset was used for training, while 20% was left aside for testing. Two stacked LSTM layers with 50 units each made up the LSTM model, which was built with Keras. To lower the chance of overfitting, dropout layers with a rate of 0.2 were added after each LSTM layer. The output layer predicted the probability distribution across bacterial species by using the softmax activation function to handle the multi-class classification job. The sparse categorical cross-entropy loss function and Adam optimiser were used to create the model, which is suitable for multi-class classification problems using integer-encoded labels. To avoid overfitting, early stopping was used to track the validation loss and terminate the training process when the validation loss stopped improving. With a batch size of 32, the model was trained for a maximum of 120 epochs. When the model’s performance was assessed on the test set after training, it showed a high degree of accuracy. The 10 most common bacterial species in the dataset were determined using predictions produced by the LSTM model. These sequence identities (sseqid) were mapped to the names of the respective bacterial species using the NCBI Entrez API, giving the computational results a biological context [19-20]. The distribution of these dominating bacterial species was displayed using bar plots, which showed their frequency, and pie charts, which showed their relative proportions throughout the dataset. This extensive research provided new insights into the microbial composition of BC tissues, finding crucial bacterial species that may be linked to the disease.

### 2.3. Breast cancer risk assessment

A total of 205 women aged 18 to 70 years, residing in the India rural areas of Morena, Sagar, Chhindwara, Gwalior, Dhar, and Betul, regions of Madhya Pradesh, were recruited. Participants who were pregnant, lactating, undergoing medication, smokers, or with a known history of chronic diseases were excluded to eliminate confounding factors. The study was approved by the Institutional Ethics Committee. From all subjects we have collected the informed written consent, and detailed sociodemographic and health-related information was collected using a structured questionnaire. Blood samples were drawn via venipuncture into EDTA-coated tubes for plasma separation. The plasma levels of granulocyte-colony stimulating factor (G-CSF), alpha-fetoprotein (AFP), and carcinoembryonic antigen (CEA) which are clinically relevant biomarkers commonly used for breast cancer risk assessment and monitoring due to their association with tumor development and progression, were measured using a standard ELISA protocol, involving biotinylated antibodies, Streptavidin-HRP conjugate, TMB substrate, and absorbance reading at 450 nm. Among the participants, 182 individuals exhibit elevated biomarker expression levels, with G-CSF ranging from 57.84 to 222.62 pg/mL, CEA from 130.62 to 383.91 pg/mL, and AFP from 8.66 to 46.93 pg/mL. These samples were subsequently selected for detailed investigation of the circulating microbiome using a developed biosensor in laboratory test settings to validate its utility for future point-of-care risk-assessment applications. The study focused on developing a detection platform to identify circulating cell-free bacterial 16S rRNA fragments found in plasma-derived circulated microbial biomarkers (CMBs). The analytical challenge in this study was not the detection of whole bacteria, but the highly sensitive and selective identification of extremely low-abundance circulating bacterial RNA fragments within complex plasma-derived nucleic acid backgrounds.

### 2.4. Circulating cell-free 16S rRNA Profiling

Cell-free circulating RNA (ccf-RNA) was extracted using the QIAMP circulating nucleic acid isolation kit. Standard venepuncture techniques were used to collect the blood samples, and the plasma was used for the further application. The extracted ccf-RNA was quantified using the Thermo Scientific™ NanoDrop™ 2000/2000c Spectrophotometer. Following quantification, complementary DNA (cDNA) synthesis was carried out using the PrimeScript 1st strand cDNA synthesis kit. The appropriate primers derived from the prepared cDNAs were used for the amplification of the 16S rRNA strand. For the amplification Qiagen RotorGene Q RT-PCR system was used, following to the specific annealing temperatures.

### 2.5. Preparation of Biotinylated Oligonucleotide Capture Probes

The 16S rRNA targeting complementary oligonucleotide probes were procured and biotinylated at the 3′ end using the Pierce™ RNA 3′ End Biotinylation Kit. For the process, 10 pmol of the oligonucleotide probes were first denatured by heating at 85 °C for 5 minutes, followed by rapid cooling to promote proper annealing. The cooled probes were then mixed with the 30% polyethylene glycol (PEG), biotinylated cytidine (bis)phosphate, 10X RNA ligation buffer containing RNase inhibitor, and T4 RNA ligase enzyme. The reaction mixture was incubated overnight at 4 °C to achieve efficient biotinylation. After incubation, nuclease-free water (NFW) was added to remove RNA ligase. To separate the aqueous phase the solution was extracted with chloroform/isoamyl alcohol (24:1). Following vigorous vortexing and high-speed centrifugation, the aqueous phase was carefully collected and combined with ice-cold 100% ethanol, glycogen and 5 M sodium chloride (NaCl). This mixture was incubated at −20 °C for one hour to facilitate precipitation of the biotinylated probes. The precipitated pellets were recovered, then washed with the 70% ethanol, and air-dried to remove residual solvent. Finally, the pellets were resuspended in NFW and stored at −80 °C until further use [20-21].

### 2.6. Preparation of Streptavidin-Coated PQDs and Biotinylated Probe Conjugation

Before the formation of the nanohybrid, carbodiimide chemistry was used to activate the carboxyl groups present on the surface of PQDs to start its coupling with streptavidin. Then a solution of EDC (1 mg/mL) and NHS (1 mg/mL) was prepared and mixed with a PQD suspension (10 mg/mL) in 1X phosphate-buffered saline (PBS), streptavidin was added to the activated PQDs at a fixed mass ratio of 1:5 (PQDs: streptavidin). The mixture was incubated at RT for around 1 hour to activate the surface of the PQDs. Then PQDs were incubated with streptavidin for a further 20 minutes at RT to increase conjugation. To enhance the efficiency of the amine-carboxyl coupling reaction, another aliquot of EDC and NHS was added to the streptavidin-PQDs conjugate mixture. This treatment optimized the conjugation efficiency, allowing for a more effective binding between the streptavidin and the activated PQDs. The mixture was stirred at RT for another hour to ensure thorough interaction between the components. Once the reaction was complete, streptavidin excess was removed from the streptavidin-coated PQDs using centrifugation, leveraging size-based separation. The supernatant was discarded, yielding a pellet of streptavidin-coated PQDs, which was subsequently stored at 4 °C for future applications. Before application the streptavidin coated PQDs were dispersed in PBS (pH 7.4) using vortex to for the uniform mixing. Biotinylated probes were then added to this dispersion in a 1:2 ratio (streptavidin-attached PQDs: biotinylated probes). The conjugation mixture was vortexed for 1 hour at RT to increase efficient streptavidin–biotin binding. Subsequently, the mixture was incubated overnight at 4 °C to allow sufficient time for the stable formation of the streptavidin–biotin conjugation. The resulting nanohybrid was used for the detection of 16S rRNA in biological samples [22-23].

### 2.7. Hydrodynamic Size Measurement

The hydrodynamic diameter of native and streptavidin-functionalized PQDs was measured by dynamic light scattering (DLS) using a Zetasizer Nano series instrument (Malvern Panalytical, UK). Measurements were performed at 25 °C in disposable polystyrene cuvettes. The PQD were dispersed with ultrapure deionized water to obtain suitable scattering intensity and to minimize multiple scattering. The samples were gently vortexed and allowed to equilibrate at 25 °C for 5 min to ensure homogeneous dispersion. Each sample was analyzed in triplicate, and the mean particle size distribution was reported. Measurements were carried out at a 173° backscattering angle, and particle size distributions were derived from the intensity autocorrelation function using cumulant and non-negative least squares (NNLS) analysis provided by the instrument software [14].

### 2.8. Co-localization analysis

Using an Olympus high-resolution fluorescence microscope, we evaluated the performance of our developed fluorescence-based nanohybrid in specifically detecting 16S rRNA amidst circulating cell-free DNA. Initially, one set of slides was prepared with streptavidin-coated PQDs, which were examined for their fluorescence intensity. In parallel, another set of slides contained a mixture of samples comprising 16S rRNA targets along with the complete nanohybrid consisting of streptavidin-coated PQDs conjugated with biotinylated oligonucleotides. To enhance detection specificity, propidium iodide (PI) was introduced as an intercalating dye within this nanohybrid, acting as a probe for the targeted PCR product of 16S rRNA. Glass slides were employed for sample preparation, ensuring optimal conditions. Each ready sample was carefully covered with a cover slip on the glass slide to protect it and maintain optimal conditions. Fluorescence imaging was performed using filter sets corresponding to DAPI (excitation ∼358 nm, emission ∼461 nm) for PQDs and TRITC/PI (excitation ∼535 nm, emission ∼617 nm). Co-localization of PQD and PI signals was confirmed by merged images, where overlapping fluorescence appeared as a combined signal, indicating successful hybridization and probe-target binding [14, 21].

### 2.9. Applicability Experiments for 16S rRNA Detection Using Fluorometry

Applicability studies of fluorescence-based detection of 16S rRNA by the developed nanohybrid system was carried out on a Spark multimode microplate reader (TECAN, Seestrasse 103, Männedorf, Switzerland). PBS was used as the negative control in all experiments. To evaluate the specificity of the streptavidin-conjugated PQDs for 16S rRNA detection, different sample combinations were used. In the control setup, a solution of PQDs with PBS was prepared and analyzed to create the fluorescence baseline. To control any changes in fluorescence intensity resulting from the streptavidin conjugation process, PQDs in conjugation with streptavidin were loaded into a separate well positioned next to the control. The important experimental step is to estimate the probe-target interaction by incorporating the 16S rRNA PCR product into the nanohybrid system with PI, an intercalating dye. The appropriate volume of the probe-target mixture was added to the next well for analysis. After an incubation period of 15 minutes, the microplate was placed in the multimode reader to measure fluorescence intensity. Using the advanced fluorescence analysis abilities of the reader, both the specificity of 16S rRNA detection and the interactions among the PI dye, streptavidin-conjugated PQDs, and the target molecules were checked [24].

### 2.10. Sensitivity Test for 16S rRNA Detection Using Nanohybrid System

To identify the lowest concentration of 16S rRNA that could be detected by the developed nanohybrid the serial dilutions (1 µg to 0.1 fg) of the 16S rRNA PCR product were prepared. PBS was used as a blank control for background fluorescence. In the experiment, the nanohybrid having streptavidin-conjugated PQDs and PI was mixed with different concentrations of the 16S rRNA PCR product in separate wells. Following the addition of the samples, the microplate was incubated for 10 to 15 minutes to allow sufficient interaction between the target RNA and the nanohybrid components. After incubation, the plate was placed into the Spark multimode microplate reader, where fluorescence intensity was recorded for each well. Data analysis, processing and visualisation were performed using Spark Control Magellan software, with a customised protocol developed for interpreting the fluorescence results. The sensitivity assay provides the detection potential of the nanohybrid system, confirming its capability to accurately identify 16S rRNA even at very low concentrations [19].

### 2.11. Specificity Investigation for 16S rRNA Detection Using Nanohybrid System

The specificity of the developed nanohybrid system for 16S rRNA detection was assessed by testing its ability to discriminate between 16S rRNA and other non-target sequences. To control background fluorescence, PBS was used. In the control setup, one well was filled with PBS containing PQDs, while another well had streptavidin-tethered PQDs suspended in PBS. To evaluate the nanohybrid’s specificity, samples containing sequences other than 16S rRNA were added to separate wells, allowing for comparison with the target RNA detection. The microplate was incubated for 10 to 15 minutes to facilitate optimal interaction between the nanohybrid and the various RNA sequences, and fluorescence intensity was measured.

## 3. Results and Discussion

The developed LSTM model addresses the inadequacy of conventional methods for analyzing noisy circulating microbiota data from SRA databases. These approaches rely on static thresholding of individual alignment outputs, which causes significant signal loss or misclassification, especially for low-abundance species crucial for risk profiling. However, our distinctive design utilizes stacked LSTM layers to concurrently integrate and holistically analyze the entire multivariate pattern of interdependent quality metrics (Figure 3). The model trained on robust instructions indicates true microbial presence by transforming taxonomic assignment from a reactive “hit-counting” method that utilizes rigid rules into a high-confidence and non-linear predictive framework [25]. This learned capability filters noise and facilitates the identification of BC-associated bacterial species with greater classification precision. This is validated by the micro-average ROC-AUC of 0.99 achieved in multiclass classification (Figure 1D), which is crucial for subsequent translational target selection.

**Figure 1:**
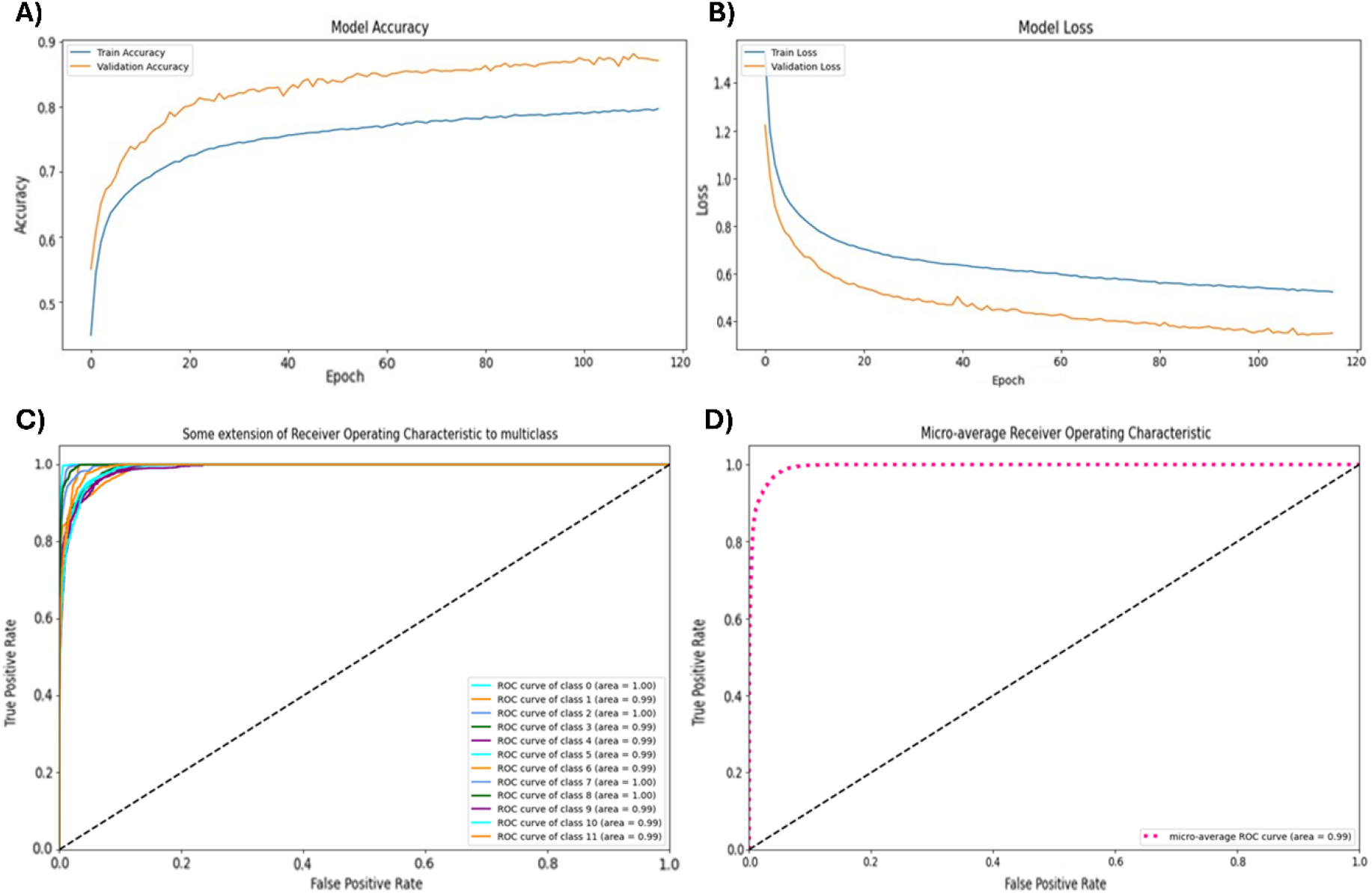
Performance and evaluation of the LSTM model predicting bacterial species. (A) Training and validation accuracy across epochs. (B) Training and validation loss across epochs during the training process. (C) ROC curves for classes 0 to 11 with AUC values near 1.00, indicating high model accuracy for each class. (D) Micro-average ROC curve with an AUC of 0.99, summarizing overall performance across all classes.

Following the optimization of the distinctive design rationale described above, the developed LSTM model exhibited robust predictive performance. During the training phase, the model achieved an accuracy of 88.16% with a corresponding loss of 0.3419, demonstrating the architecture’s efficacy in effectively classifying bacterial species from the complex metagenomic dataset. This high level of performance validates the model’s capability to transform standardized BLAST-derived taxonomic annotations into meaningful predictive patterns. Crucially, Machine Learning analysis successfully learned the complex rules within critical metrics such as sequence identity, alignment length, and E-value. When applied to unseen data, the LSTM model maintained high predictive stability, achieving a validation accuracy of 87.18% and a validation loss of 0.3483. This negligible reduction in performance compared to the training set confirms the framework’s stability and robust generalization capability **(Figure 1A and 1B)**.

In classified 11 distinct classes of the dataset, the performance of the LSTM model was evaluated using the micro-average Receiver Operating Characteristic (ROC) curve **(Figure 1C)**. The ROC curve plots the True Positive Rate (TPR) in contrast to the False Positive Rate (FPR) at different threshold settings, offering a visual representation of the trade-off between sensitivity and specificity. The Area Under the Curve (AUC) value of 0.99 was achieved, indicating exceptional performance. An AUC close to 1 reflects a strong ability to differentiate between negative and positive classes, whereas the diagonal reference line (AUC = 0.5) corresponds to the performance of a random classifier with no predictive capability **(Figure 1D)**. These findings highlight the exceptional robustness and accuracy of the LSTM model, with the ROC curve serving as a valuable tool for evaluating its performance in multiclass classification tasks. The high AUC value further emphasizes the model’s reliability in making accurate predictions for the bacterial species in the dataset.

The model identified four bacterial species as the most prevalent within the breast cancer tissue samples: *NZ_LM831024*.*1* (*Pseudomonas aeruginosa)* (19.4%), *NZ_LT906445*.*1 (Veillonella parvula)* (19.3%), *NZ_CP045110*.*1* (*Acinetobacter baumannii)* (16.8%), and *NZ_LR134275*.*1* (*Streptococcus vestibulacis)* (8.5%) (**Figure 2A)**. These findings indicate an association of these microbial species with the breast cancer tissue microbiome and suggest their potential involvement in disease-related biological processes. To further visualize the distribution, a bar chart titled “Top 10 Bacterial Species Distribution” was created, illustrating the frequency of the top 10 bacterial species identified in the dataset. The chart revealed that *NZ_LM831024*.*1* had the highest frequency, slightly exceeding 30,000, followed by *NZ_LT906445*.*1*, with a frequency just below 30,000. Other species, such as *NZ_CP045110*.*1* (approximately 25,000), *NZ_LR134275*.*1* (around 15,000), and *NZ_CP086333*.*1* (slightly below 15,000), were also prominent in the dataset **(Figure 2B)**. These top 10 species reflect the microbial composition of the tissue samples, underscoring their prevalence and potential impact on the pathophysiology of breast cancer. The integration of BLAST-based taxonomic profiling and LSTM-based AI modeling provides a powerful framework for uncovering the microbial landscape of cancer tissues and contributes to advancing the understanding of the role of microbiota in disease development.

**Figure 2:**
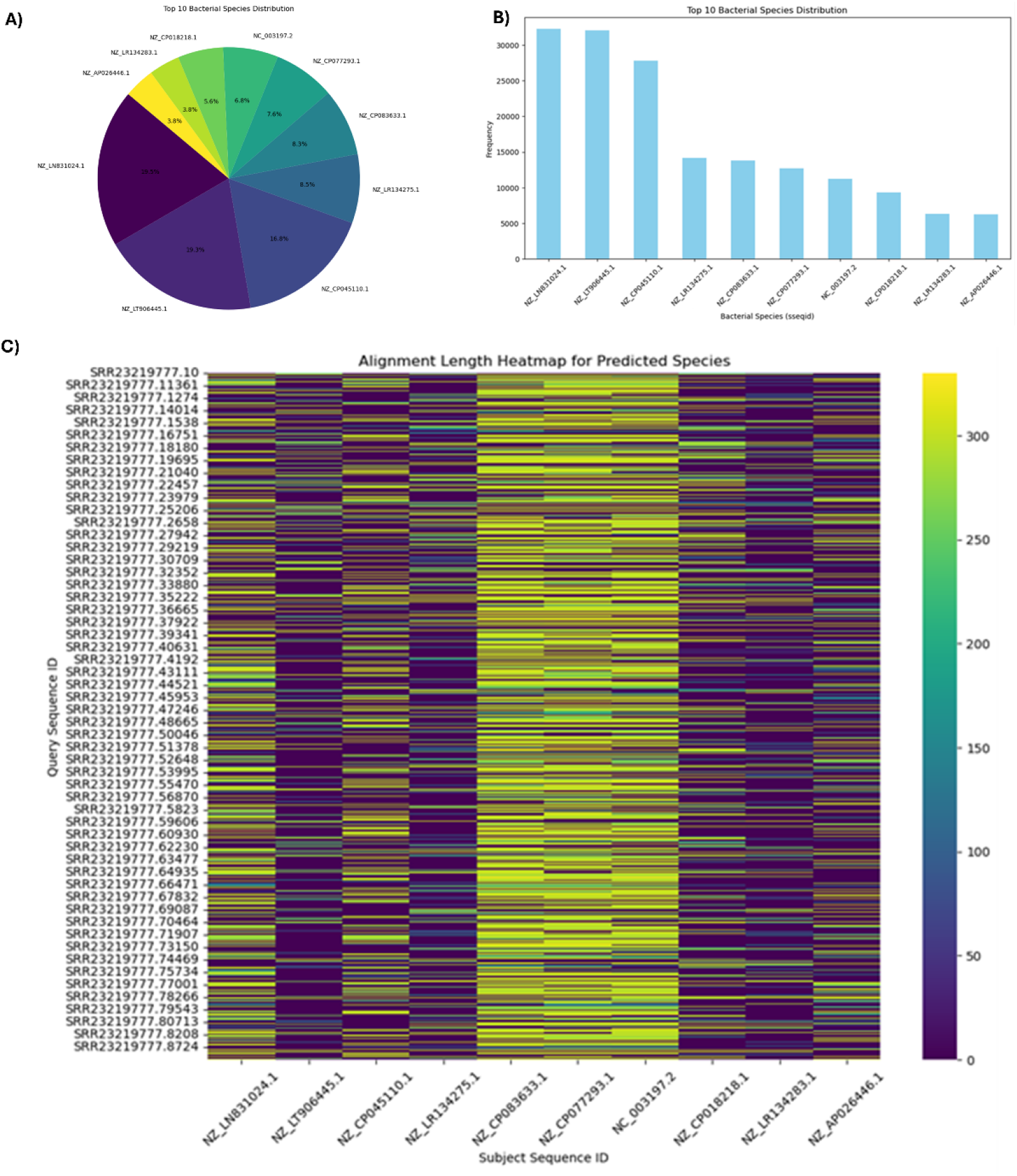
Distribution and alignment of the ten most abundant bacterial species predicted by the LSTM model. (A) Percentage distribution of the predicted bacterial species. (B) Frequency distribution of the predicted bacterial species. (C) Heatmap displaying the alignment lengths between the different predicted bacterial species.

**Figure 3:**
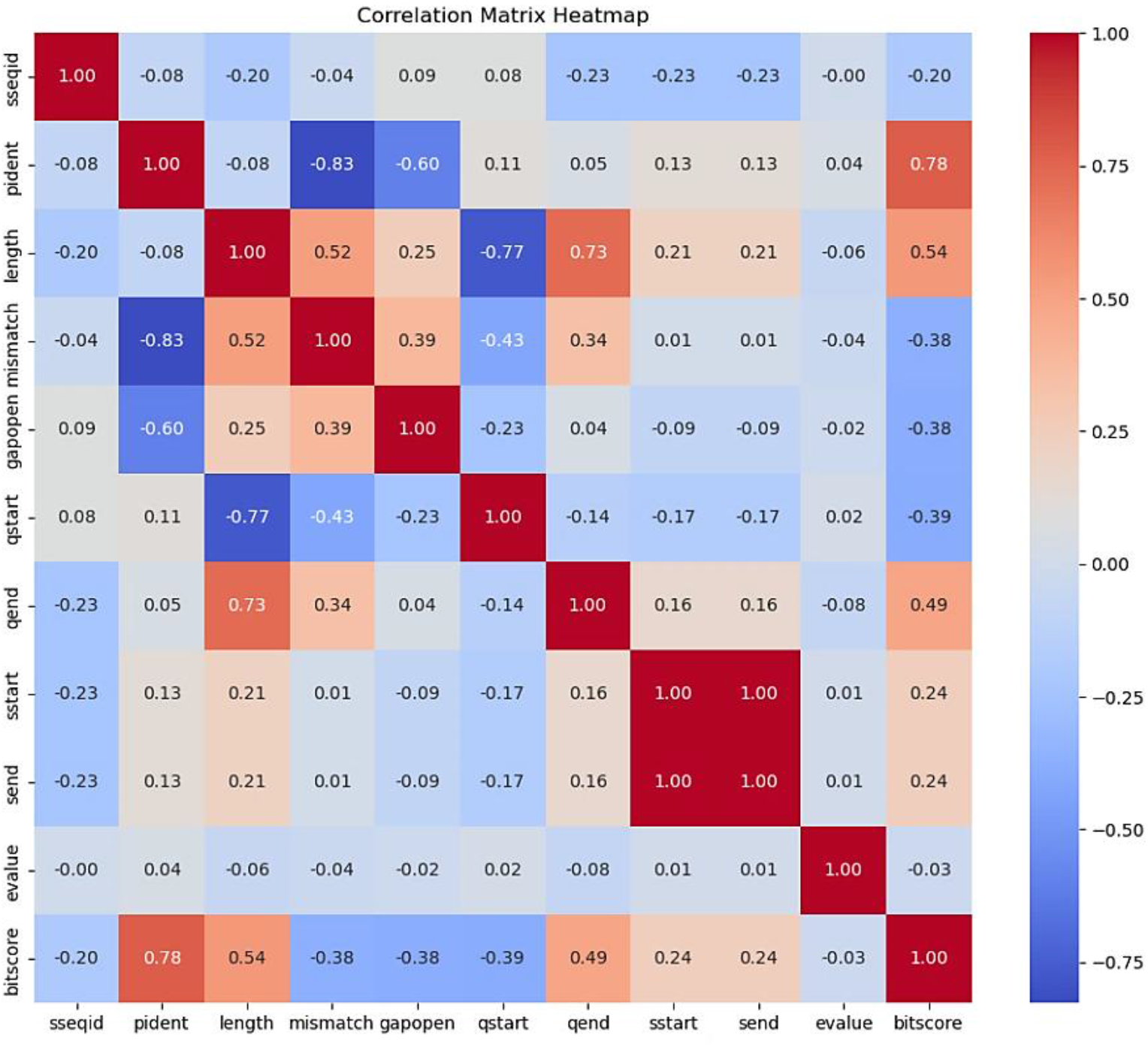
The heatmap illustrates a correlation matrix, showing relationships between features like sseqid, pident, length, and others. It uses colors ranging from dark blue (strong negative correlation, - 1) to dark red (strong positive correlation, 1), with white indicating no correlation (0).

The “Alignment Length Heatmap for Predicted Species” visually represents the alignment lengths between query sequences and reference subject sequences. The heatmap uses a color gradient, with darker colors indicating shorter alignments and lighter colors signifying longer alignments. The Y-axis displays query sequences identified by SRR numbers, while the X-axis lists subject sequences by their reference identifiers. The heatmap highlights clusters where longer alignments suggest stronger relationships between specific query and subject sequences. This tool effectively identifies trends and patterns in sequence alignments, aiding in genomic data interpretation and supporting further research **(Figure 2C)**.

The correlation matrix, which is often represented in a heatmap, visually captures the relationships between different features in the dataset. The degree to which two features are correlated are showed by each value in the matrix. The value close to 1 represents strong positive correlation, and the values close to -1 show a strong negative correlation. By analyzing this matrix, one can identify pairs of features that are highly correlated, providing valuable insights, how these features influence each other. Features that are strongly correlated may contain redundant information, suggesting that one could be removed to simplify the model without much loss of predictive power. On the contrary, features with low or negative correlation may represent independent or complementary information, which could be important for making distinct predictions. The correlation matrix helps to refine the feature selection process, improving model performance by focusing on the most relevant and non-redundant features **(Figure 3)**.

Following the AI model prediction for bacterial species, PQDs were used for the fabrication of nanobiosensors for detecting 16S rRNA in collected samples. PQDs have recently attracted significant attention owing to their unique optical and electrical properties, which make them highly suitable for advanced nanosensor development and nanomedicine applications. Their advantages in nucleic acid biomarker detection include high photoluminescence quantum yields, size-dependent optical behaviour, and tuneable emission wavelengths. Because of these characteristics, PQD-based nanosensors may detect new nucleic acid biomarkers, such as circulating DNA, miRNA, lncRNA, and other nucleic acids that are suggestive of infectious and cancerous disorders, with extraordinary sensitivity and selectivity [15, 20]. PQDs integration into biosensing systems enables quick, real-time detection, which helps with early diagnosis, particularly when low-abundance nucleic acids may be difficult to detect using conventional diagnostic techniques. Conjugating PQD-based nanosensors with complementary nucleic acid probes enables targeted hybridisation with sequences in nanomedicine. PQDs’ optical characteristics may alter in observable ways as a result of this process, offering a very sensitive platform for measuring target at incredibly low concentrations [11].

The PQD nanohybrid was constructed using 4 sets of streptavidin-coated PQDs and biotinylated oligonucleotide probes with sequence complementary to the 16S rRNA of 4 selected proteobacteria for selective binding. In this study, an extensive RT-PCR panel screening was carried out to assess the expression profiles of three selected Proteobacteria 16S rRNA sequences previously linked to pathogenic mechanisms involved in BC progression. The objective was to identify microbial signatures that may contribute to BC risk modulation. (**Figure 4**) presents the relative expression levels of these bacterial 16S rRNA sequences in high-risk BC samples compared with low-risk samples, using human mitochondrial DNA (mtDNA) 16S rRNA as the internal reference for normalisation. The analysis revealed elevated expression levels of the four targeted bacterial species, *P. aeruginosa, V. parvula, A. baumanii, and S. vestibulacis* in samples categorized as high-risk for BC. These findings highlight the potential involvement of these bacterial species in the underlying molecular and cellular mechanisms that contribute to BC progression. The choice of human mtDNA as a reference control provided a robust normalization framework, ensuring accurate quantification of microbial RNA expression across varied sample types [23]. Furthermore, this differential expression emphasizes the relevance of circulating microbiota as potential biomarkers for assessing BC risk. The observed patterns underscore the critical need for advanced molecular diagnostics, such as the integration of nanobiosensor-based approaches.

**Figure 4:**
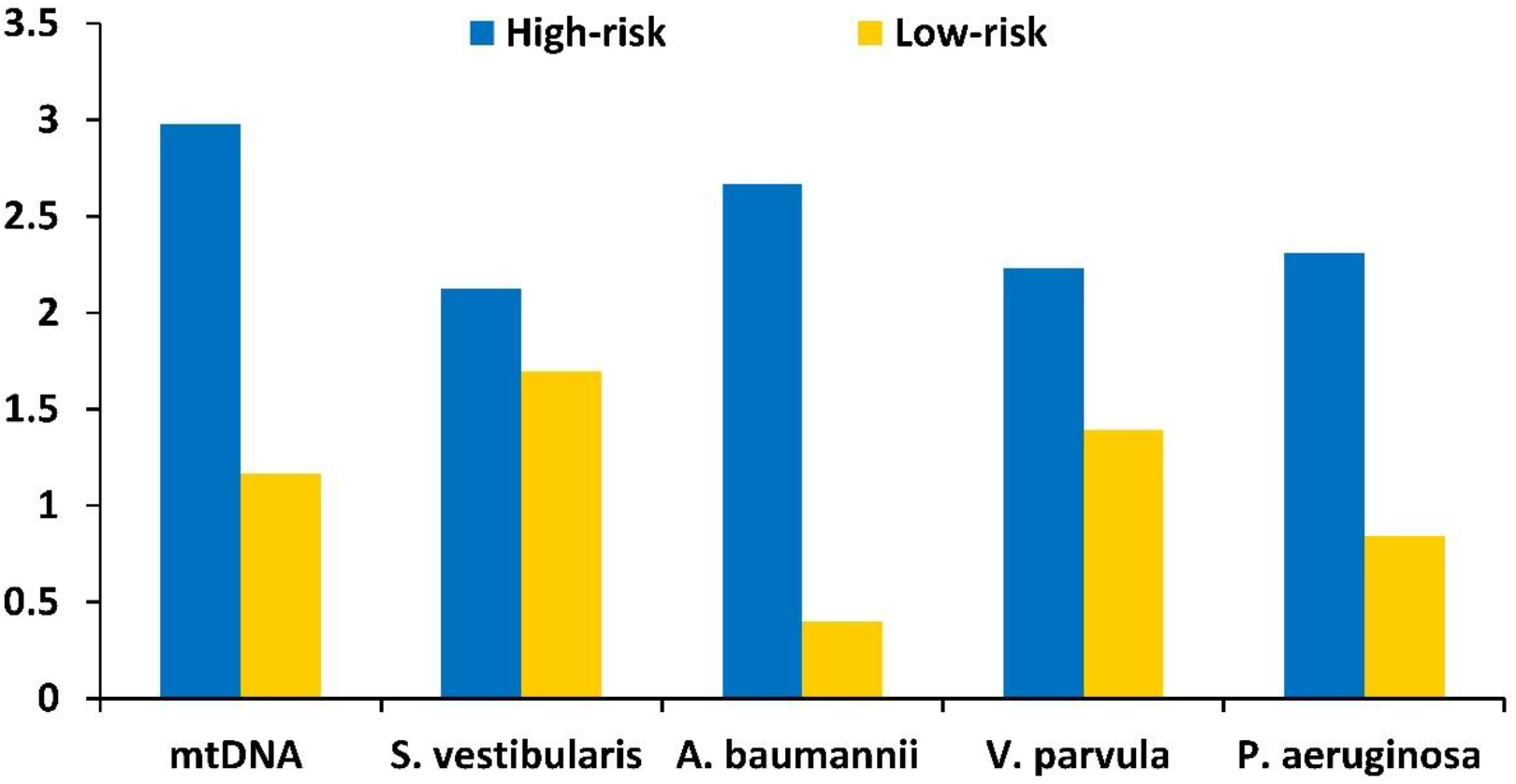
Graph showing the ΔΔCt values of 16S rRNA for S. vestibularis, A. baumannii, V. parvula, and P. aeruginosa.

After fabricating the 4 nanohybrids, the DLS measurement indicates that the Native PQDs **(Figure 5A)** exhibited a narrow and monomodal size distribution in the 20-40 nanometers range, indicating a well-dispersed nanoparticle population with good colloidal stability. Whereas, streptavidin-functionalized PQDs **(Figure 5B)** have a noticeable shift toward larger hydrodynamic diameters with a broader size distribution extending into the 80-150 nm and 300-500 nm submicron ranges. The increase in particle size is attributed to the formation of a streptavidin layer on the PQD surface and possible protein-mediated interparticle interactions. The observed increase in hydrodynamic diameter confirms the successful surface functionalization of PQDs with streptavidin, enabling their use in biotin-streptavidin-mediated biosensing applications.

**Figure 5:**
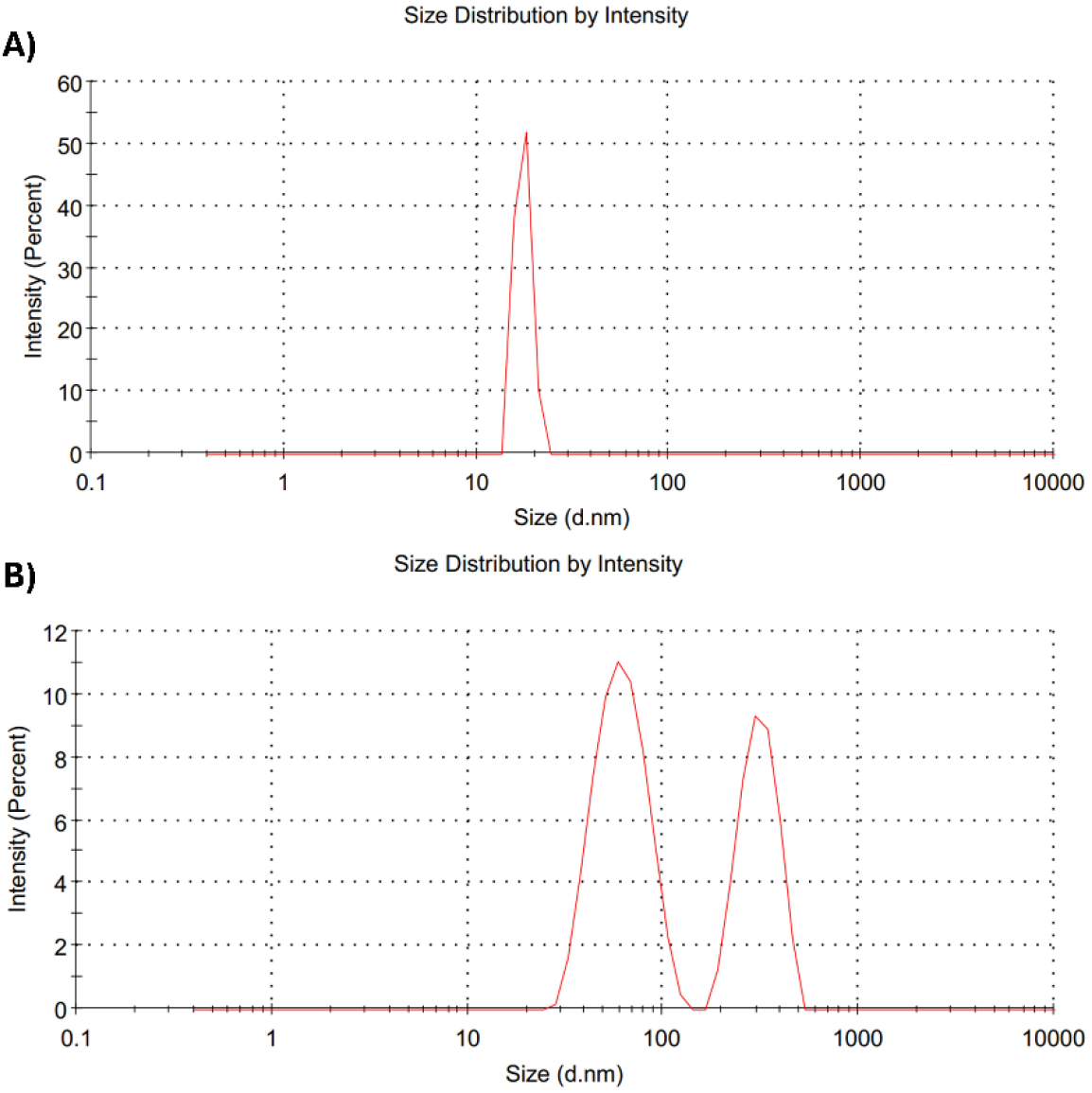
Size distribution of PQDs: (A) indicate a native PQDs showing a narrow distribution and (B) is the streptavidin-functionalized PQDs exhibiting increased hydrodynamic diameter, indicating successful surface functionalization.

After the size estimation, we undertook a comprehensive evaluation of their fluorescence properties and detection capabilities through the use of microscopy and flow cytometry, as illustrated in **Figure 6**. To investigate the native fluorescence characteristics of the PQDs, we analyzed the nanohybrids under DAPI, employing Propidium Iodide (PI) as an intercalating dye to visualize the binding interactions. The results of the fluorescence microscopic evaluation of the conjugation are presented in Panel I of **Figure 6**. In the initial image (Image A), we observe a striking blue fluorescence that indicates the presence of streptavidin-conjugated PQDs, indicate the successful incorporation into the nanohybrid structure. The subsequent image (Image B) reveals an exciting development: after introducing the biotinylated probe, we detected specific target bacterial 16S rRNA sequences from organisms including *P. aeruginosa, V. parvula, A. baumanii* and *S. vestibulacis*. This detection is characterized by a vivid red fluorescence emission, attributable to the intercalation of PI. The binding of PI takes place along with the formation of a double-stranded complex between the complementary probe and the target 16S rRNA by inserting itself into the major and minor grooves of the double helix, thereby confirming the probe’s specificity for the target gene.

**Figure 6:**
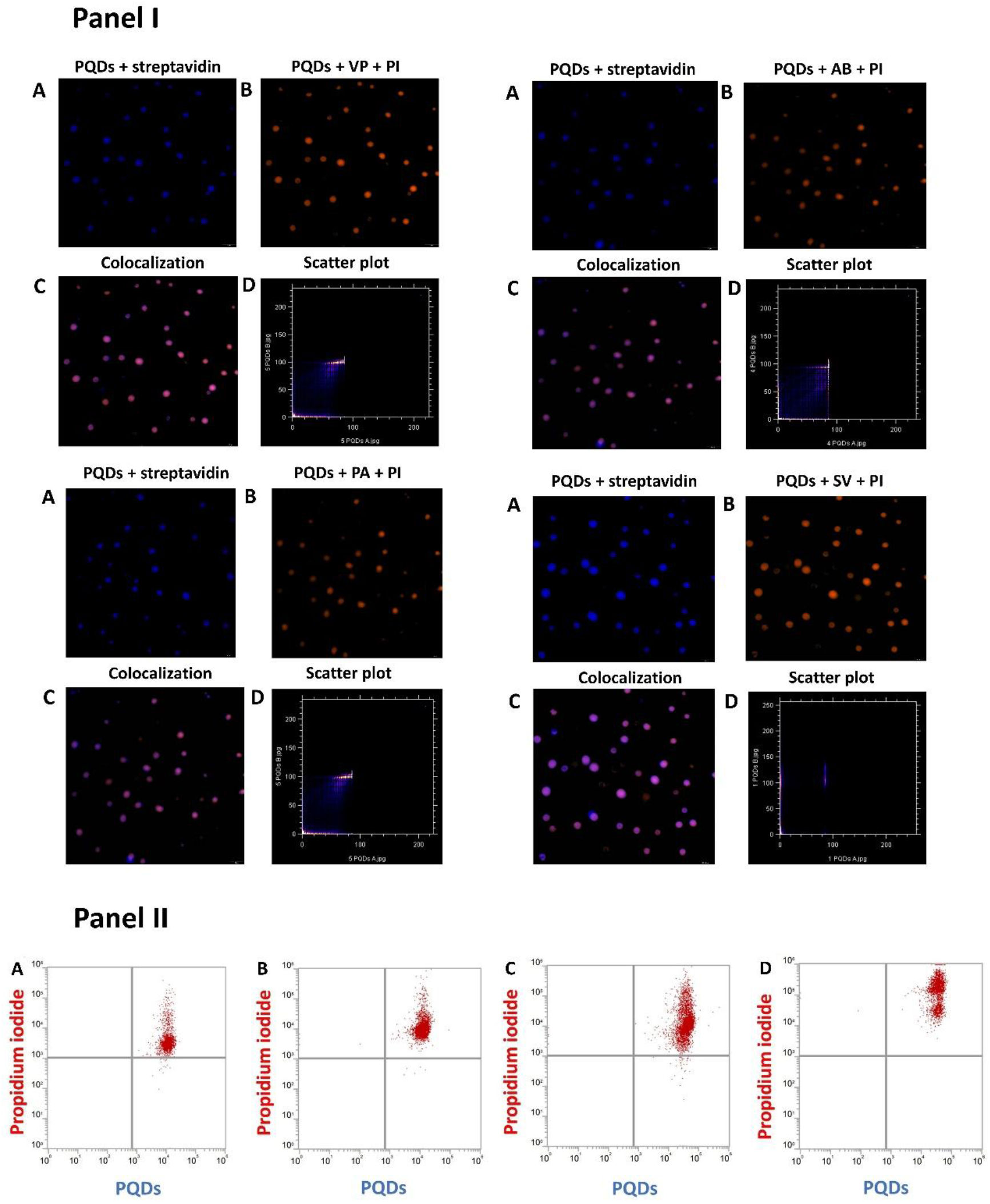
Validation of the nanohybrid system for bacterial 16S rRNA detection. * Panel I (Microscopy): (A) Intrinsic PQD fluorescence. (B) PI dye fluorescence. (C) Merged image showing colocalization (purple), confirming specific 16S rRNA binding. (D) Scatter plot of spatial correlation between PQD and PI signals. Panel II (Cytometry): Dot plot demonstrating simultaneous 16S rRNA detection of four target bacterial species (P. aeruginosa, V. parvula, A. baumannii, S. vestibulacis). The PI fluorescence shift confirms successful probe hybridization.

Figure 6, Panel I provides a detailed evaluation of the synthesized nanoanalytical framework for the detection of bacterial 16S rRNA, utilizing high-resolution fluorescence microscopy and flow cytometry. In Panel I, Image (A) demonstrates the intrinsic fluorescence emitted by the perovskite quantum dots (PQDs), (B) displays the fluorescence signal of the intercalating dye, PI, Image (C) presents the colocalization of fluorescence signals from PQDs and PI, with the merged signals appearing as purple fluorescence. This overlap tells the successful conjugation and specific binding of the nanohybrid system to the target bacterial 16S rRNA. The specificity of this interaction is further supported by the scatter plot shown in Image (D), which represents the spatial correlation between the PQD and PI signals. Panel II indicates a dot plot from nano cytometric analysis, demonstrating the simultaneous detection of the 16S rRNA from all four bacterial species in circulating microbial biomarkers (CMB). The upper-right quadrant of the dot plot corresponds to the fluorescence-positive population of PI, indicating successful hybridization of the biotinylated probe with 16S rRNA [26]. This interaction confirmed the selective binding of the probe to the 16S rRNA of target bacterial species and provided the foundation for subsequent experiments. In image C, a careful examination of images (A) and (B) reveals an intriguing phenomenon: the overlapping fluorescence signals converge to create a striking purple colour. This vivid hue displays the successful formation of the nanohybrid. In addition, image D provides a detailed scatter plot that offers a visual representation of the colocalization, making it easier to interpret the relationship between the different signals. To measure its effectiveness in detecting the target 16S rRNA in biological samples, additional evaluations were conducted using nano cytometry to deepen our understanding of the method’s capabilities. The results, shown in Panel II of Figure 6, display a pronounced shift in the fluorescence of PI, suggesting a robust interaction when the 16S rRNA hybridized with specific bacterial oligonucleotide probes embedded within the nanohybrid structure. This shift indicates effective intercalation of PI within the double-stranded arrangement of the oligonucleotide probes, capturing the specific 16S rRNA of notable bacteria, including *P. aeruginosa, V. parvula, A. baumannii*, and *S. vestibulacis*, all within the intricately designed nanoframework.

The practical applicability of streptavidin-conjugated PQD for the detection of 16S rRNA underwent a comprehensive evaluation, yielding significant insights into the behaviour of the nanohybrid system. The experimental analysis showcased a pronounced reduction in fluorescence intensity when PQDs bound to streptavidin, observed through the blue filter. This decrease in fluorescence is likely due to the disturbing of PQD surface in COOH-NH_2_ coupling conditions. The attachment of biomolecules to the PQD surface can disturb the system’s delicate energy balance and produce defects.

It is interesting to mention that the fluorescence intensity of streptavidin modified PQDs was significantly amplified upon the addition of biotinylated probes, suggesting an efficient interaction within the system. The combination of PI and the biotinylated probe produced the highest fluorescence intensity, as determined by further investigations performed under a PI filter (excitation ∼535 nm, emission ∼617 nm). These findings highlight the nanohybrid system’s great potential for the accurate detection of 16S rRNA **(Figure 7 (Applicability)**.

**Figure 7:**
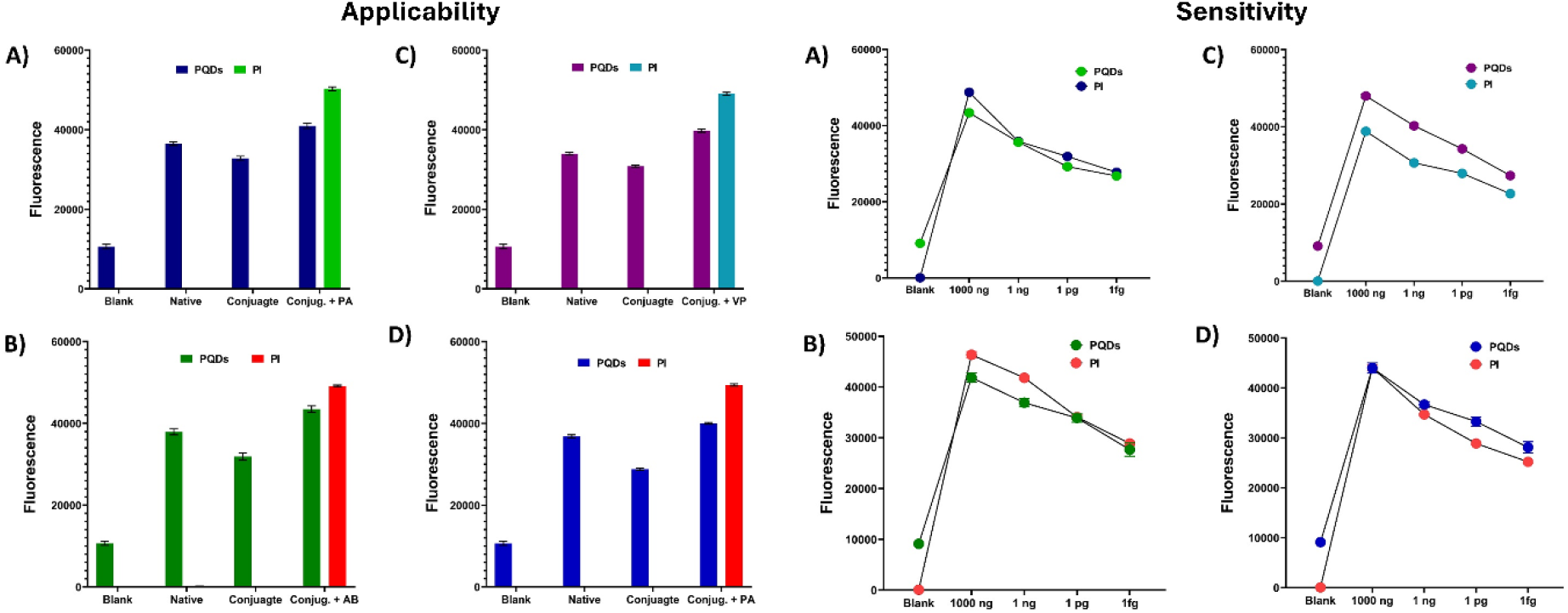
Performance of the fabricated nanohybrid for bacterial 16S rRNA detection. (I) Applicability: Graphs illustrating the applicability of the nanohybrid for detecting 16S rRNA in (A) Streptococcus vestibularis, (B) Acinetobacter baumannii, (C) Veillonella parvula, and (D) Pseudomonas aeruginosa. (II) Sensitivity: Graphs depicting the sensitivity of the developed nanohybrid for detecting 16S rRNA in (A) Streptococcus vestibularis, (B) Acinetobacter baumannii, (C) Veillonella parvula, and (D) Pseudomonas aeruginosa.

The sensitivity analysis reveals the detection threshold of the nanohybrid system for 16S rRNA. The results display a significant analytical sensitivity with the detection of bacterial 16S rRNA concentrations as low as **1 fg**. This level of sensitivity highlights the potential clinical applicability of the nanohybrid system for ultra-sensitive detection of microbial biomarkers in complex biological matrices. The detection sensitivity was found to correlate near to linear with fluorescence intensity and target concentration. Maximum fluorescence intensity was obtained at an optimal target concentration of **1000 ng**, ensuring efficient signal transduction and reliable detection. However, a progressive decline in sensitivity was noted with decreasing concentrations of 16S rRNA. Despite this decline, the system maintained a robust limit of detection (LOD), remaining highly sensitive up to 20.58 fg, 15.1 fg, 20.9 fg, 20.1 fg for *P. aeruginosa, A. baumannii, V. parvula*, and *S. vestibulacis respectively*, which is attributed to the enhanced binding efficiency between the target and the biotinylated probe within the nanohybrid framework **(Figure 7 (Sensitivity)**.

To evaluate the ability of the nanohybrid system to reliably distinguish target genes within complex biological samples, detection of bacterial 16S rRNA sequences in the presence of diverse nonspecific components was performed. The analysis involved two different 16S rRNA fragments, one of which was human mitochondrial 16S rRNA (mt-16S rRNA), known to share substantial sequence similarity with bacterial 16S rRNA. As shown in Figure 10, the nanohybrid displayed markedly higher binding efficiency for bacterial 16S rRNA compared to mt-16S rRNA. This selective binding is primarily attributed to the complementary sequence specificity of the biotinylated oligonucleotide probe, which preferentially hybridizes with bacterial 16S rRNA. Additionally, the smaller base-pair size and structural differences in bacterial 16S rRNA further enhance its binding affinity with the nanohybrid system, as highlighted in **Figure 8**. The preferential interaction underscores the nanohybrid’s capacity to discriminate between closely related sequences, even in the presence of high concentrations of non-specific nucleic acids.

**Figure 8:**
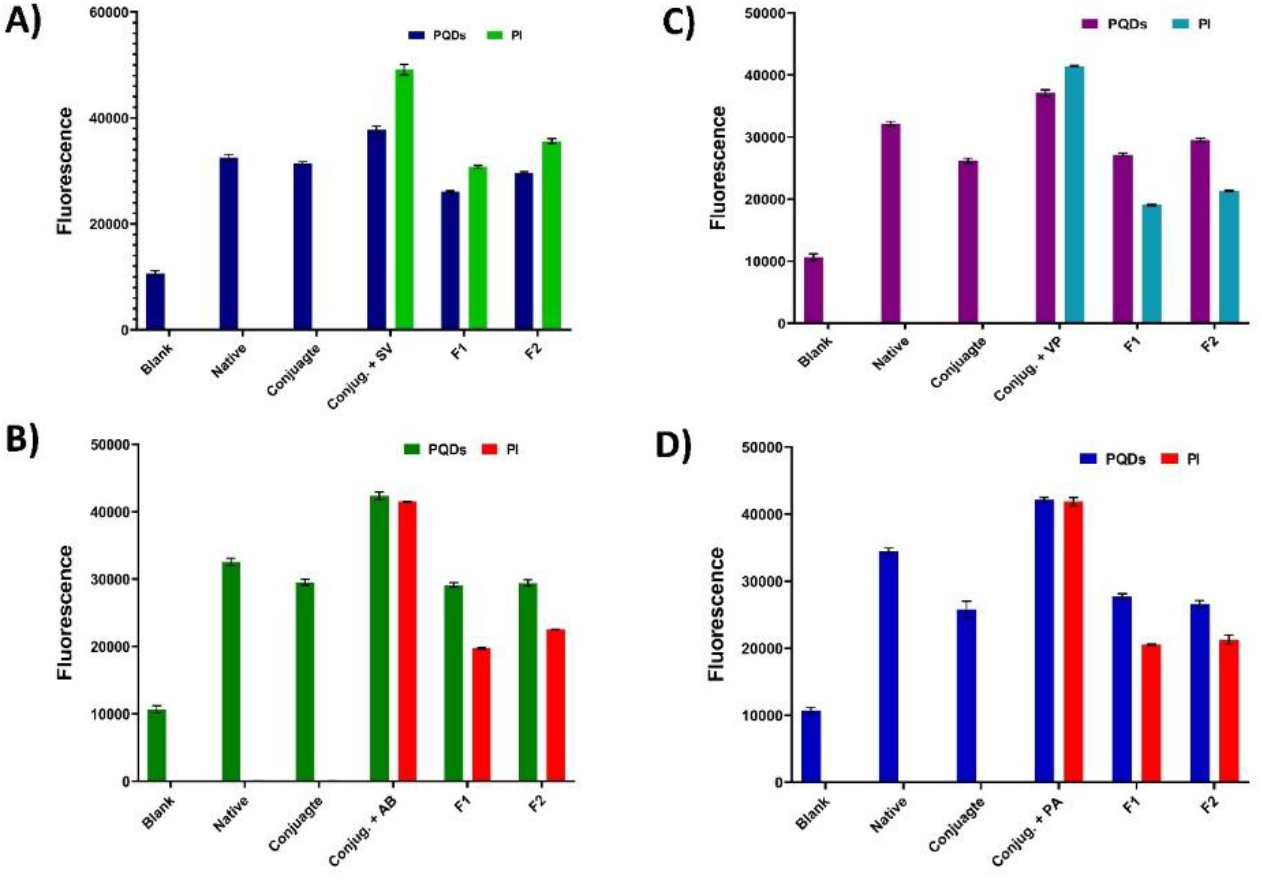
Graph depicting the specificity of the developed nanohybrid for detecting 16S rRNA in (A) Streptococcus vestibularis, (B) Acinetobacter baumannii, (C) Veillonella parvula and (D) Pseudomonas aeruginosa.

**Figure 9:**
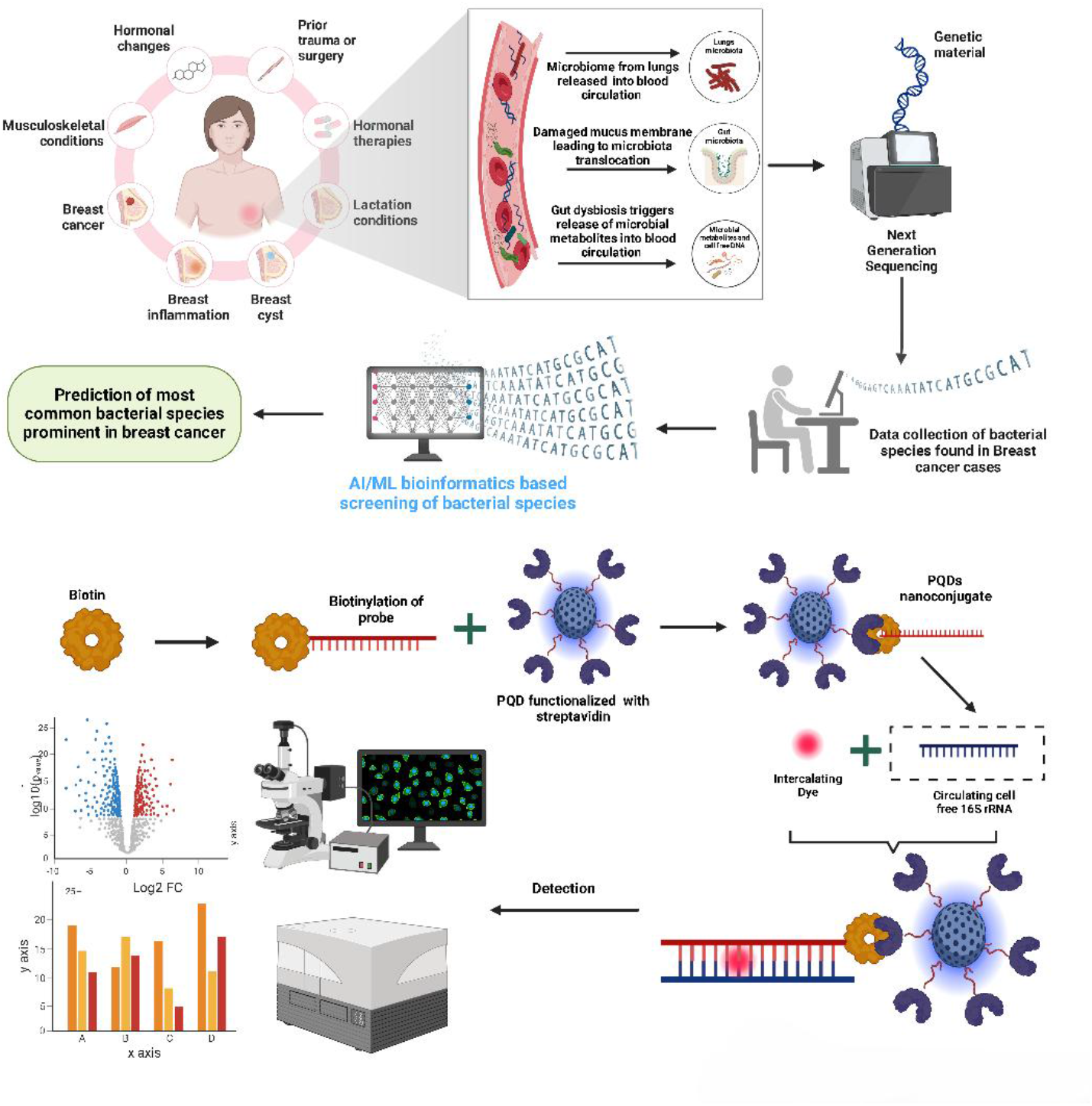
An illustration of a proposed method for identifying and detecting bacterial species that contribute to breast cancer: samples from individuals with different risk factors are subjected to AI/ML-based bioinformatics and next-generation sequencing, in which bacterial DNA is biotinylated and captured by streptavidin-conjugated PQD. The data obtained, which are then examined for differential abundance, aid in diagnosis and possible treatments.

The specificity of the nanohybrid system is attributed to its design, which integrates PQDs conjugated with streptavidin and biotinylated oligonucleotide probes. The strong affinity of the biotin-streptavidin interaction ensures nanohybride stability, while the oligonucleotide probe’s high sequence fidelity minimizes cross-reactivity with non-target sequences. Furthermore, the intercalation of PI into the double-stranded complex formed during hybridization amplifies the fluorescence signal, providing a reliable readout for target recognition. This high specificity emphasizes the nanohybrid system’s utility for the precise detection of CMBs in complex biological fluids. Its ability to selectively target bacterial 16S rRNA, while excluding closely related sequences such as mt-16S rRNA, validates its potential as a robust diagnostic tool. Future work will focus on extending the specificity study to additional microbial species and testing the nanohybrid system across diverse biological matrices. This will further validate its potential for integration into point-of-care diagnostic platforms, providing a precise and efficient solution for microbial biomarker detection in clinical and environmental settings.

The earlier developed sensors such as electrochemical immunosensors are often restricted by its reliance on sophisticated equipment and ability to monitor circulating microbial nucleic acid biomarkers [27-29]. While, other nanotechnology based sensors lack specific integration with microbial profiling and unbiased AI-assisted computational disease prediction strategy [30, 31]. The nanohybrid system developed herein overcome these limitations by integrating the enhanced photoluminescence efficiency coupled with precise targeting of cancer-related microbial nucleic acids and advanced computerized prediction ability. In addition, the combination of novel 16S rRNA biomarker, simplified optical processing, and enhanced classification accuracy through computational integration make it suitable for advanced clinical biosensing.

## 4. Conclusion & translational perspective

In conclusion, the present study showcased a robust fluorescence-based nanohybrid system that utilize PQDs, streptavidin, and biotinylated oligonucleotide probes for the rapid and accurate identification of CMBs in BC. This study introduces a fluorescence-based nanohybrid system that enables rapid, accurate identification of CMBs in clinical samples. By integrating PQDs with streptavidin and biotinylated oligonucleotide probes, the system offers high specificity and sensitivity for detecting microbial species associated with BC. The practical applicability of the devised nanohybrid technology was assessed utilising plasma-derived circulating microbial RNA samples under physiologically relevant background conditions. Investigations into specificity conducted in the presence of non-target nucleic acids, such as human mitochondrial 16S rRNA, revealed no nonspecific fluorescence interference, hence affirming the platform’s durability in intricate biological matrices. It attained femtogram-level sensitivity with fast fluorescence detection with brief incubation times. In contrast to traditional molecular diagnostic methods like RT-PCR and next-generation sequencing, which typically require extensive sample preparation, expensive equipment, and lengthy analysis times, the current PQD-based fluorescence nanoplatform offers benefits such as accelerated analysis, improved fluorescence sensitivity, a simpler workflow, and potential adaptability for point-of-care diagnostic applications. This robust fluorescence-based platform represents a significant step forward in BC microbiome research, providing rapid detection of microbial biomarkers and insights into their roles in cancer development. Future work will focus on validating this platform across diverse clinical samples and integrating it into POC diagnostic systems for greater accessibility and impact in resource-limited settings.

## Acknowledgment

The authors thank the technical staff member, Mr. Vinay Singh Raghuvanshi, for his assistance in collecting samples.

## Ethical Declaration

The study received approval from the Institutional Ethics Committee (IEC) of ICMR-NIREH under the reference number Project ID 5/3/8/1/GIA-ITR, IRIS Cell ICMR ID No.: 2020-9466.

## Author Contribution

PKM devised the concept, developed the methodology, and supervised the experiments; NN, PR and VG performed most experiments; VG, IYG characterized samples; and VG, AB, SRL designed the figures; VG and OG performed the data analysis and interpretation; OG, RT, IYG reviewed the manuscript; and NN, VG, AB and PKM drafted the original manuscript.

## Data Availability

The data supporting this study’s findings are available upon request from the corresponding author.

